# A death pheromone, oleic acid, triggers hygienic behavior in honey bees (*Apis mellifera L.*)

**DOI:** 10.1101/231902

**Authors:** Alison McAfee, Abigail Chapman, Immacolata Iovinella, Ylonna Gallagher-Kurtzke, Troy F. Collins, Heather Higo, Lufiani L. Madilao, Paolo Pelosi, Leonard J. Foster

## Abstract

Eusocial insects live in teeming societies with thousands of their kin. In this crowded environment, workers combat disease by removing or burying their dead or diseased nestmates. For honey bees, we found that hygienic brood-removal behavior is triggered by two odorants – β-ocimene and oleic acid – which are released from brood upon freeze-killing. β-ocimene is a co-opted pheromone that normally signals larval food-begging, whereas oleic acid is a conserved necromone across arthropod taxa. Interestingly, the odorant blend can induce hygienic behavior more consistently than either odorant alone. We suggest that the volatile β-ocimene flags hygienic workers’ attention, while oleic acid is the death cue, triggering removal. Bees with high hygienicity detect and remove brood with these odorants faster than bees with low hygienicity, and both molecules are strong ligands for hygienic behavior-associated odorant binding proteins (OBP16 and OBP18). Odorants that induce low levels of hygienic behavior, however, are weak ligands for these OBPs. We are therefore beginning to paint a picture of the molecular mechanism behind this complex behavior, using odorants associated with freeze-killed brood as a model.

## Introduction

Disease and parasite transmission is a constant threat in dense insect societies^1-3^. Ants^4-8^, termites^9-11^, and honey bees^12-16^ have evolved social mechanisms of disease resistance which mitigate this risk and improve the collective health of their colonies. Ants transport dead nestmates to their midden heaps, termites bury or entomb their dead in graves, and honey bees remove dead and diseased brood from the hive. E. O. Wilson described these processes as ‘necrophoresis,’^4^ or the movement of dead individuals away from the colony. Necrophoresis reduces pathogen reservoirs, inhibiting the spread of diseases and parasites from fallen nestmates to those who endure^1,2,4^.

In honey bees (*Apis mellifera*), one dominant form of necrophoresis is hygienic behavior^13,14^. Hygienic honey bee workers will identify and remove diseased, dead, and sometimes parasitized larvae, prepupae, and pupae from the colony. This is an effective defense against major diseases, including chalkbrood (*Ascosphaera apis*)^17,18^, American foulbrood (*Paenibacillus larvae*)^14,19^, and the devastating *Varroa* mite (*Varroa destructor*)^13,14,20,21^. When highly hygienic colonies are challenged with these pests and pathogens, they are less likely to develop clinical symptoms than non-hygienic hives, and are more likely to recover and survive^19,22,23^.

The underlying mechanism of the behavior has only been partially deciphered. Like other social insects, honey bees identify their diseased nestmates via chemical cues^24-27^; however, since partway through development (late 5^th^ instar larvae and older) the brood becomes capped and completes development in the confines of a sealed wax cell, the workers have an added challenge. The physical barrier between the bees who execute the behavior and the brood interferes with their ability to detect their targets. Detecting the dead, diseased, or parasitized capped brood is thought to rely on volatile odorant signals that permeate the wax cell cap^26^, but very few hygienic behavior-inducing odorants have been identified and confirmed behaviorally^27,28^. Swanson *et al.*^27^ found that a volatile chalkbrood odorant (phenethyl acetate) was a strong hygienic behavior-inducer, and Nazzi *et al.* showed that a volatile *Varroa*-associated odorant ((Z)-6-pentadecene) does the same^28^. Non-volatile cues have not yet been investigated behaviorally in honey bees, despite including some of the most taxonomically conserved necrophoretic and necrophobic compounds (e.g. oleic acid and linoleic acid)^1,6,9,10,29-34^.

Hygienic honey bees have superior olfactory sensitivity compared to non-hygienic honey bees^24-27^, which likely depends in part on differences in antennal gene expression^23,35-38^. In a search for antennal biomarkers for hygienic behavior, we previously identified two odorant binding proteins – OBP16 and OBP18 – that significantly correlated with colony hygienic score^35^. Antennae are honey bees’ main olfactory appendages, and OBPs aid odorant signal detection by binding and transporting hydrophobic odorant molecules from the antennal pores to the olfactory nerves^39^. Despite some tantalizing inferences, OBP16 and 18 have not been mechanistically linked to hygienic behavior.

Previously, we compared odorant profiles of freeze-killed pupae and healthy pupae to find candidate hygienic behavior-inducing compounds^40^. Although freeze-killing is not a natural means of death, it is a relevant system because the freeze-killed brood assay^41^ is the main method for determining colonies’ level of hygiene. We identified two new candidate compounds that were significantly more abundant in freeze-killed brood: oleic acid and β-ocimene. Oleic acid is a non-volatile, oily substance which acts as a death cue in eusocial and non-eusocial insects^1,6,9,10,29-34^. For example, oleic acid stimulates necrophoretic behavior in multiple ant species^6,31,32^, as well as termites^9,10^. In isopods^29^, caterpillars^29^, crickets^30^, cockroaches^34^, and bumble bees^33^, it induces avoidance behavior, presumably as a mechanism to avoid the risk associated with disease or predation indicated by other dead insects. β-ocimene, on the other hand, is a volatile honey bee brood pheromone that is normally a larval food-begging signal^42^. β-ocimene emitted from larvae is also known to inhibit worker ovary development^43,44^, regulate the nurse-to-forager transition^44^, and stimulate foragers to forage^45,46^. The queen also produces β-ocimene, which also contributes to inhibiting worker egg-laying^47^. Prior to our work in 2017^40^, β-ocimene and oleic acid had not been linked to hygienic behavior in honey bees.

In the present work, we investigate oleic acid and β-ocimene’s roles in hygienic behavior using behavioral assays, electrophysiology, and OBP ligand binding assays. Our behavioral assay overcomes a major hurdle in testing the hygienic behavior-inducing capacity of odorants: by adding odorants through the resealable cells of Jenter™ queen cages, we can add individual odorants to brood cells while maintaining perfect integrity of the wax cell walls and cap. By this method, we show that even the non-volatile oleic acid induces hygienic behavior, but the blend of β-ocimene and oleic acid induced hygienic behavior most strongly and consistently. However, we have not ruled out the possibility of a brood effect induced by odorant contact toxicity. Electroantennogram (EAG) recordings of bees from a hygienic colony show that at a hive-realistic temperature and humidity, oleic acid induces only slightly above-background antennal nerve responses, suggesting that it is only detectable upon contact or extremely close proximity. *In vitro* ligand binding assays show that both β-ocimene and oleic acid are strong ligands for at least one of OBP16 or OBP18 (which are upregulated in hygienic bees’ antennae)^23,35^, and the two odorants we tested which do not substantially induce hygienic behavior were poor ligands. Taken together, we propose a mechanistic model where the co-opted, volatile brood pheromone (β-ocimene) works together with an evolutionarily conserved death cue (oleic acid) via interactions with hygienic behavior-associated odorant binding proteins (OBP16 and OBP18) to induce hygienic behavior.

## Results

Previously, we found that β-ocimene and oleic acid were both emitted more strongly from freeze-killed brood compared to live brood, making them promising candidates as putative hygienic behavior-inducers. To test if these odorants are sufficient to induce brood removal, we developed a front-way odorant assay (**Figure 1A**), which involves uncapping patches of brood (30 cells each, in two technical replicates per colony) and dispensing 1 µl of either neat (100%) or diluted (1%) odorant standards on the brood.

**Figure 1.**
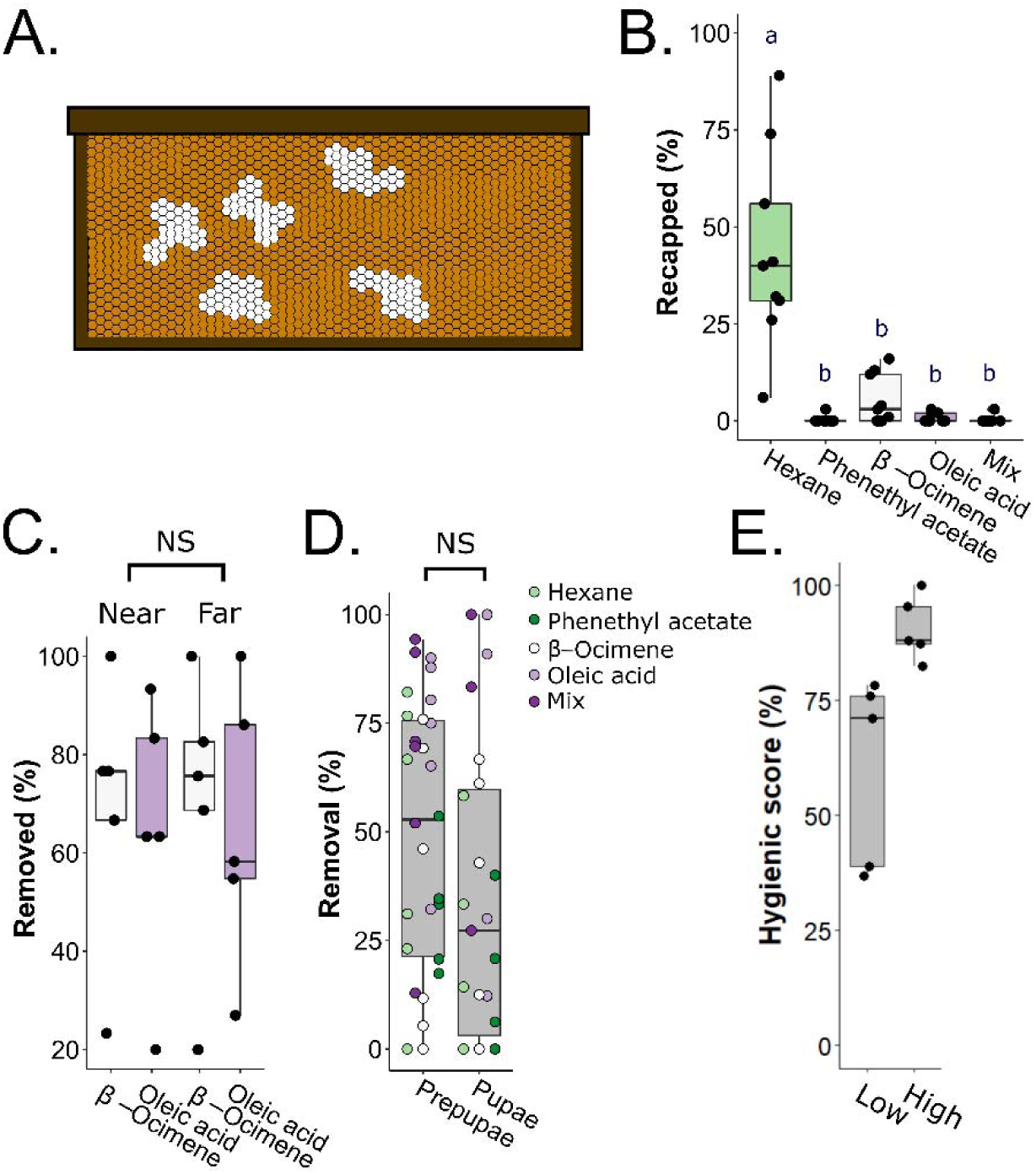
Front-way odorant assay preliminary tests. **A)** Schematic of the front-way assay. Patches of capped brood (∼30 cells in technical duplicate per colony) developing naturally in a standard frame were uncapped (white patches) and 1 µl of odorants (β-ocimene, oleic acid, a **1:1** v/v mix of the two, phenethyl acetate or hexane) at either 1% or 100% concentrations (v/v in hexane) were dispensed onto the brood. Frames were incubated in the colony’s brood box for 3 hours before recording removal rates. **B)** Post-front-way assay recapping frequencies. Data from N = 9 colonies were analyzed with a one-way ANOVA (level: odorant; F = 13.3, p = 2.4e-8) followed by a Tukey HSD test. Letters indicate groups that are significantly different from one another Tukey HSD p < 0.05). **C)** Preliminary test for a patch proximity effect. N = 5 colonies were tested, varying the distance between β-ocimene and oleic acid patches (near = patches on the same frame, separated by one band of untreated capped brood; far = patches on different frames separated by two untreated brood frames). We analyzed the data by a two-way ANOVA and found no effect of patch (F = 0.025, p = 0.88) nor interactive effect between patch and odorant (F = 0, p = 1.0). D) Preliminary test for a brood age effect. We performed the front-way assay on N = 9 colonies and calculated the percent prepupa and pupa removal. Due to variability in patch composition, not every colony had the same number of replicates for each stage and dose (see Table 1 for all sample sizes). Data were analyzed with a four-way ANOVA (levels: odorant, age, hygienicity, dose), which identified no significant effect of age nor interactions with any other factors, followed by a Tukey HSD test. 1% and 100% refer to odorant concentrations. All boxes depict the interquartile range (IQR) and the whiskers span 1.5*IQR.

First, we confirmed that hexane was an appropriate negative control by recording the recapping frequencies following the treatments (N = 9 colonies). We found that after just three hours, an average of 44% of the hexane-treated cells were recapped, which was significantly higher than for all other odorants (**Figure 1B**; one-way ANOVA followed by Tukey HSD; β-ocimene: p = 2e-7; oleic acid: p = 1e-8; mix: p = 1e-8; phenethyl acetate: p = 1e-8). The next highest was β-ocimene, with 5.4% recapped. The others all had recapping frequencies of 1% or less, indicating that the brood were no longer accepted by the workers.

**Table 1:**
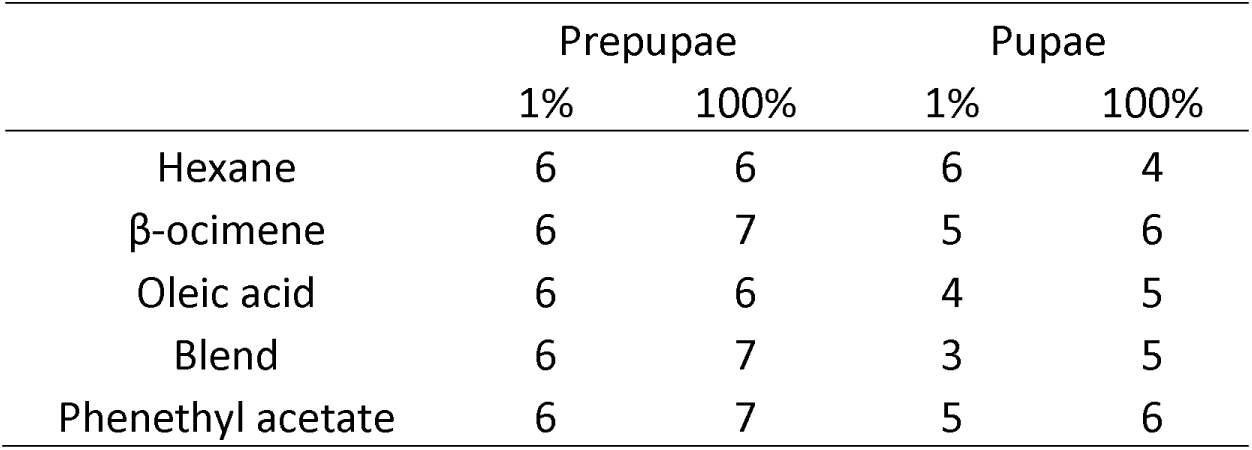
Replicate information for age-related brood removal measurements

Next, we sought to confirm that there was no effect of patch proximity on brood removal. To test this, we treated patches of ∼30 cells with β-ocimene or oleic acid, and separated the patches by either one band of untreated cells (’near’ treatments) or located the patches on two different frames, with two untreated brood frames separating them (’far’ treatments). We did this for N = 5 colonies, and found no effect of patch proximity on brood removal rates (**Figure 1C**; two-way ANOVA; levels: odorant, proximity; F = 0.025, p = 0.88). In another test, we found that workers removed treated pupae and prepupae at similar rates (**Figure 1D**; four-factor ANOVA; levels: dose, odorant, hygienicity, age; F = 0.84; p = 0.36; see Table 1 for sample sizes). Therefore, we combined data for the two ages and used the front-way assay to test if colonies with higher hygienicity responded to the odorants differently than colonies with lower hygienicity. We tested N = 5 colonies with high hygienicity (freeze-killed brood score > 80%) and N = 5 colonies with low hygienicity (freeze-killed brood score < 80%) (**Figure 2A**), and found significant effects of dose, odorant, and hygienicity (**Figure 2B**; three-factor ANOVA; dose: F = 61.2, p = 4.3e-11; odorant: F = 19.8; p = 7.1e-11; hygienicity: F = 20.2, p = 2.7e-5).

**Figure 2.**
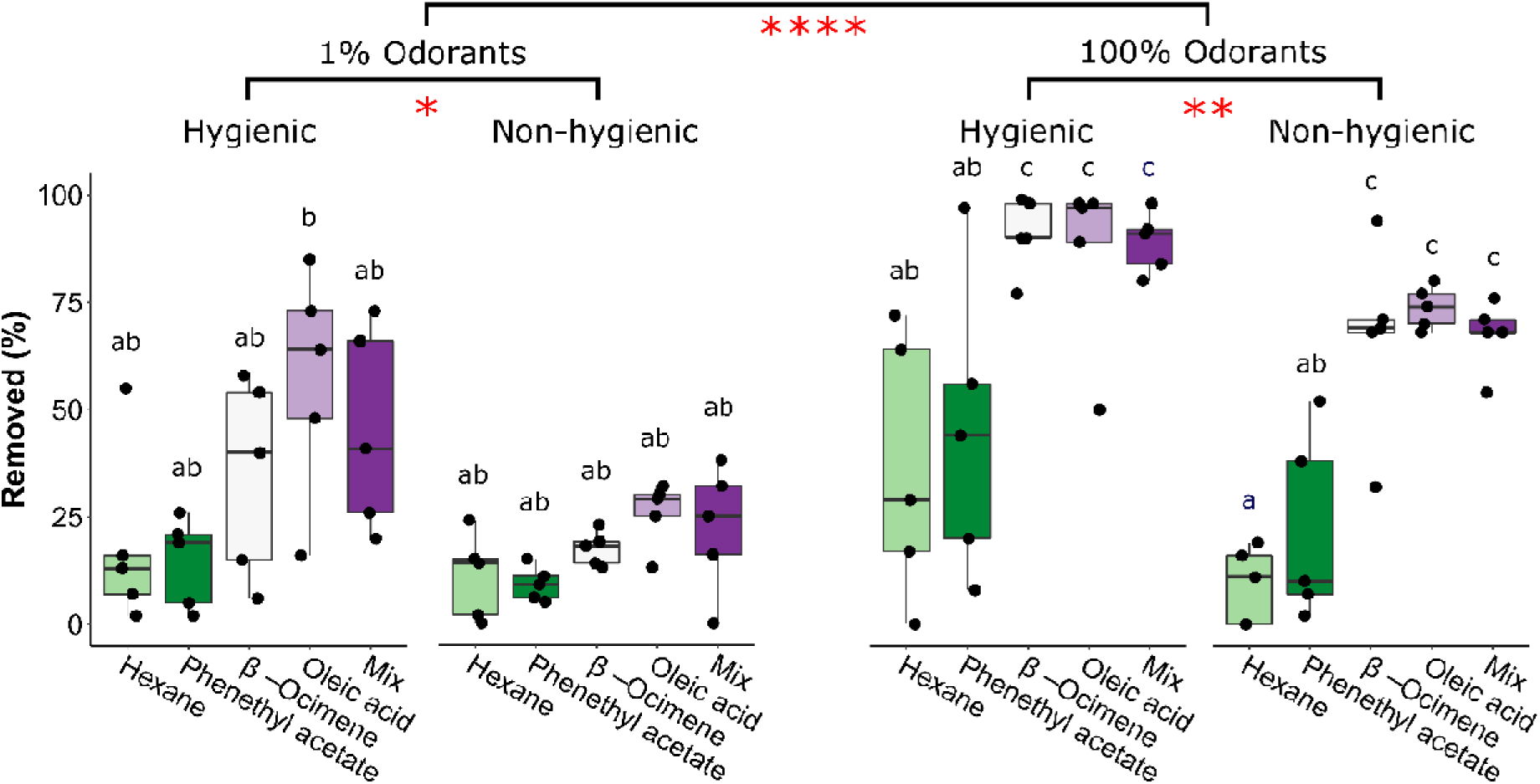
Front-way odorant assays to investigate effects of hygienicity. **A)** Distribution of hygienic scores for the tested colonies. 10 colonies were tested in total. The lowest-scoring 5 were assigned to the ‘low hygienicity’ group (scores < 80%) and the highest-scoring 5 were assigned to the ‘high hygienicity’ group (scores > 80%). **B)**. Post-front-way assay removal frequencies. Hexane is the negative control and phenethyl acetate (a chalkbrood odorant) was meant to be the positive control. Data from 5 low hygienicity and 5 high hygienicity hives were analyzed with a three-factor ANOVA (levels: dose, odorant, hygienicity; dose: F = 61.2, p = 4.3e-11; odorant: F = 19.8; p = 7.1e-11; hygienicity: F = 20.2, p = 2.7e-5), followed by a Tukey HSD post-hoc test. Significance code (Tukey HSD): * p < 0.05, ** p < 0.01, **** p < 0.0001. Boxes depict the interquartile range (IQR) and the whiskers span 1.5*IQR. Letters indicate groups that are significantly different from one another at Tukey HSD p < 0.05).

As expected, brood treated with neat odorants were removed significantly more frequently compared to those treated with diluted odorants. We had intended phenethyl acetate to be a positive control odorant, but surprisingly, we found that it induced similar brood removal as the negative control (hexane), both of which were the lowest of all those we tested. In the neat odorant treatments, β-ocimene, oleic acid and their blend all induced significantly higher brood removal relative to hexane (Tukey HSD; p = 0.0034, p = 0.0075, and p = 0.0049 respectively), but in the diluted odorant treatments, none of the odorants induced significantly different brood removal. However, their relative patterns still reflect what’s observed in the neat odorant treatments.

We expected colonies with higher hygienicity to respond more strongly to the odorant stimuli than colonies with lower hygienicity. We found that indeed, the higher hygienicity colonies removed significantly more treated brood overall in both the neat odorant treatments (Tukey HSD; p = 0.0084), as well as the diluted treatments (p = 0.011). This agrees with previous electroantennography studies showing that hygienic bees’ antennae are more sensitive to disease odorants than non-hygienic bees^24,25^.

The front-way odorant assay is a quick method of gauging if odorants can induce brood removal, but it cannot test for odorant transmission through the physical barrier of the wax cap. To investigate the odorants in a more realistic scenario, we devised a new assay using the Jenter™ system that allows us to treat brood with odorants while maintaining the integrity of the brood cells. We call this the back-way odorant assay (**Figure 3A**), since we add the odorants through the back of the brood cell. Briefly, we place a queen in a Jenter™ cage until she lays eggs in the comb of the cage, then release her and allow the workers to rear the brood until it is capped. The back of the Jenter™ cage is equipped with removable plugs that enable odorants to be added inside the cell without disturbing the delicate wax cell cap, and plugged again to close the brood cell. We used this method to add neat hexane, β-ocimene, oleic acid and the odorant blend to 9-10 brood cells each, before and after pupation (N = 5 colonies for each age). We found that after incubating in the hive for 20 h, β-ocimene did not induce significantly more brood removal relative to hexane (**Figure 3B;** two-factor ANOVA followed by Tukey HSD; p = 0.82 for pre-pupal brood and p = 0.10 for post-pupal brood). However, oleic acid strongly induced pre-pupal removal (p = 0.0004) and marginally non-significant post-pupal removal (p = 0.057). The odorant blend induced the most consistently high brood removal of them all, which was significant for both brood ages (p = 0.0004 for pre-pupal and p = 0.0003 for post-pupal).

**Figure 3.**
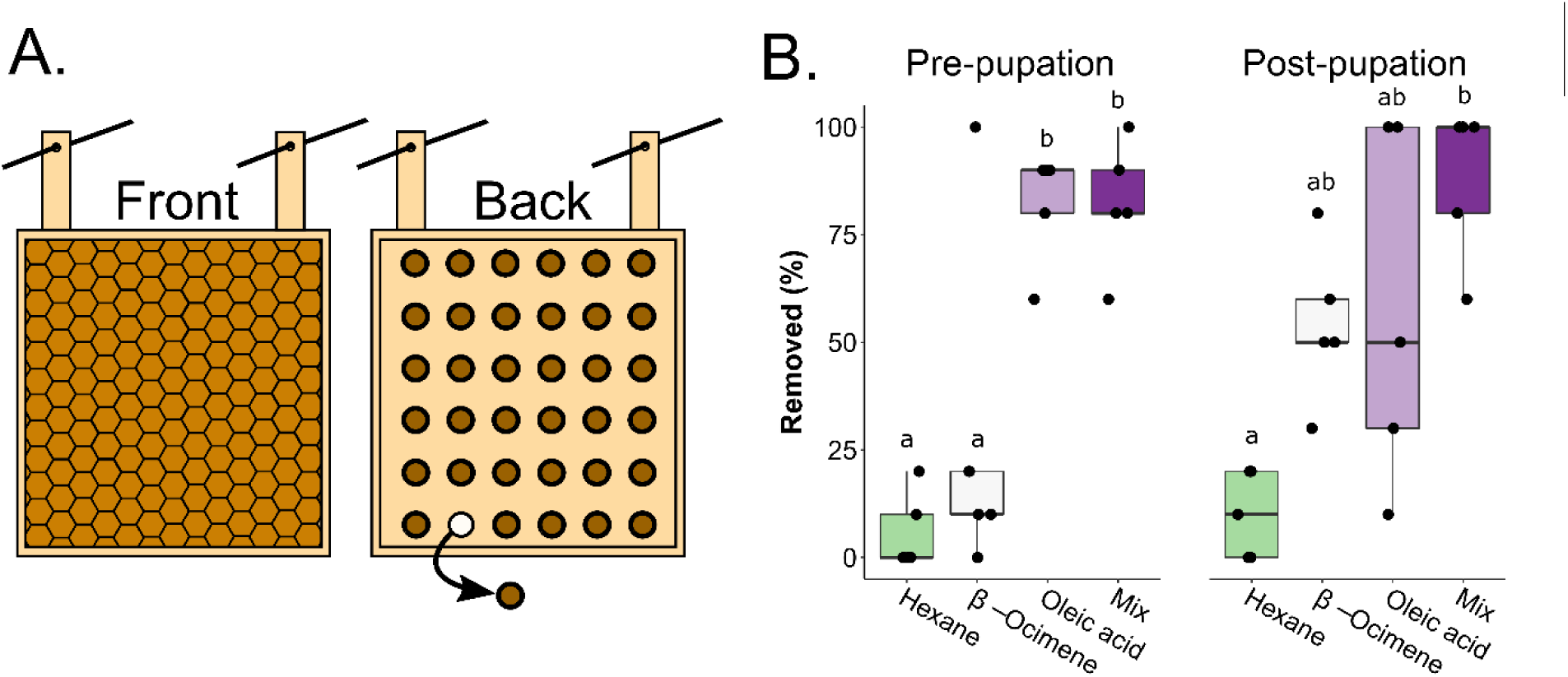
Back-way odorant addition assays. **A)** Schematic of the back-way assay. Queens were caged in a Jenter™ queen rearing cage (a hanging square of artificial comb) until she populated the cells with eggs. The queens were released and brood were allowed to develop until capping (front view). We treated brood cells with neat odorants in a semi-random design through the cell plugs (back view, brown circles), then the odorant-impregnated brood was incubated in the colony for 20 h to allow time for odorant diffusion, uncapping, and removal. Diagrams are not to scale. The actual Jenter™ cage has ∼ 100 removable plugs (one every 3^rd^ cell). **B.)** We treated pre-pupal and post-pupal brood with each odorant (9-10 brood cells for each age and odorant, N = 5 colonies). Data was analyzed using a two-factor ANOVA (levels: age and odorant) followed by a Tukey HSD post hoc test. There was a significant effect of odorant (F = 20.3, p = 1.51e-7), no significant effect of age (F = 0.16, p = 0.694), and no significant interactive effect (F = 1.9, p = 0.157). Letters indicate groups that are significantly different from one another (Tukey HSD p < 0.05). Boxes depict the interquartile range (IQR) and whiskers span 1.5*IQR.

To try to explain the patterns of pre-pupation and post-pupation brood removal, we investigated changes in the background volatile and non-volatile odorant profiles that could confound with our odorant treatments. To do this, we performed solid-phase micro-extraction gas chromatography-mass spectrometry (SPME GC-MS) on extracts from 5^th^ instar larvae, prepupae, and pupae. We analyzed N = 5 independent brood, from 5 different colonies, for each stage. We also used a hexane wash (with the same replicate structure as before) to extract cuticle compounds from these life stages and analyzed them by GC-MS as well, capturing the less volatile signals. We found that β-ocimene abundance changed most significantly according to age (one-way ANOVA, Benjamini-Hochberg corrected 1% FDR; p = 0.0010, q = 0.01), with relatively high amounts emitted in 5^th^ instar larvae and prepupae, and low amounts in pupae (**Figure 4A** and **B**). Two other minor chromatogram components were also differentially emitted (compounds 2 and 4, corresponding to isopropanol and 2-pentanone, respectively). Other volatile compound identifications are reported in **Table S1**. The hexane wash identified many branched chain hydrocarbons which were differentially emitted with age but importantly, oleic acid was not among the identified molecules for any of the three developmental stages (**Table S2**).

**Figure 4.**
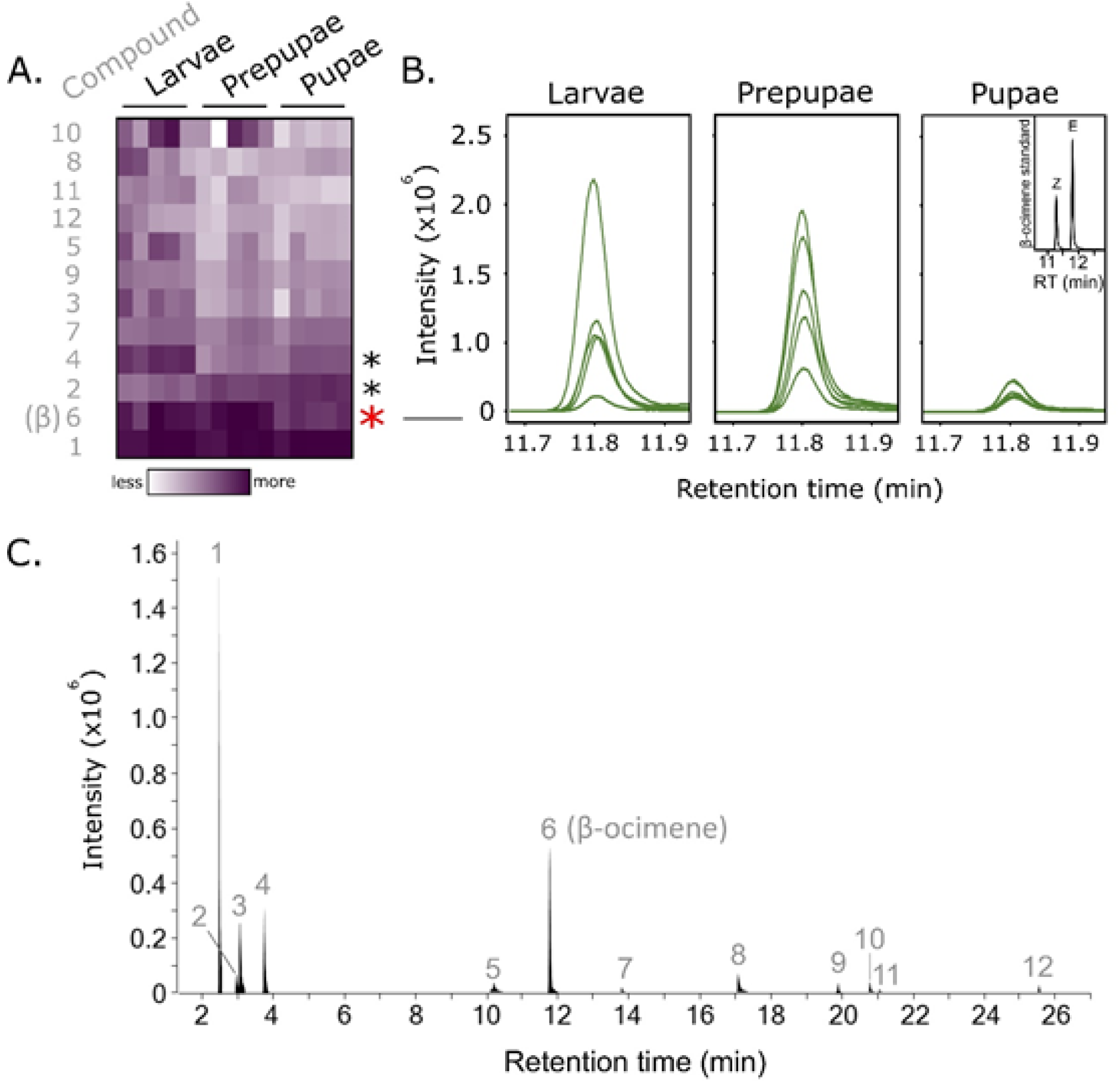
β-ocimene abundance in larvae, prepupae and pupae. We performed solid phase micro-extraction gas chromatography mass spectrometry (SPME-GC-MS) on extracts from 5^th^ instar larvae, prepupae and pupae (N = 5 colonies each). **A)** Heatmap showing intensities of all integrated peaks. Areas under the curve were compared between ages using a one-way ANOVA and Benjamini-Hochberg correction (5% FDR). Each row corresponds to peak intensities belonging to a different compound. β-ocimene, the most significantly different compound, is indicated with a red asterisk, while two other significantly different compounds (matching to isopropanol (2) and 2-pentanone (4)) are indicated with black asterisks. Raw GC-MS data is available at http://github.com/AlisonMcAfee/test. **B)** Chromatogram traces of the β-ocimene peak. Its identity was confirmed with a synthetic standard (inset chromatogram). Based on its retention time, only the E isomer was identified in the brood. **C)** Example SPME-GC-MS total ion chromatogram. Numbers correspond to compounds labelled in **3A**. Further compound identity and abundance information is available in **Table S1**.

Previously, we reported that stimulating honey bee antennae with oleic acid yielded no measurable nerve depolarization signal above the background stimulus of air alone^40^. Since we clearly observe that oleic acid can induce hygienic behavior in brood removal assays (including when the brood cell cap remains in-tact), we questioned if the workers were detecting oleic acid-treated cells by olfaction or some other sense (e.g. gustation). To investigate this further, we replicated the electroantennography experiment (N = 13 left antennae and N = 14 right antennae) comparing oleic acid to background stimulation, but at a temperature that better-matches in-hive conditions. When we administered warmed oleic acid (at approximately 33°C) we found that it stimulates worker antennae only slightly more than blank stimuli (**Figure 5**). There was also a significant effect of odorant (two-way ANOVA; levels: odorant, side; F = 12.4; p = 2.3e-5), with β-ocimene and the odorant blend inducing significantly higher antennal nerve depolarizations than oleic acid in left antennae (p = 0.011 and p = 0.016, respectively). The same comparisons yielded a marginally non-significant response in the right antennae (p = 0.085 and p = 0.086, respectively).

**Figure 5.**
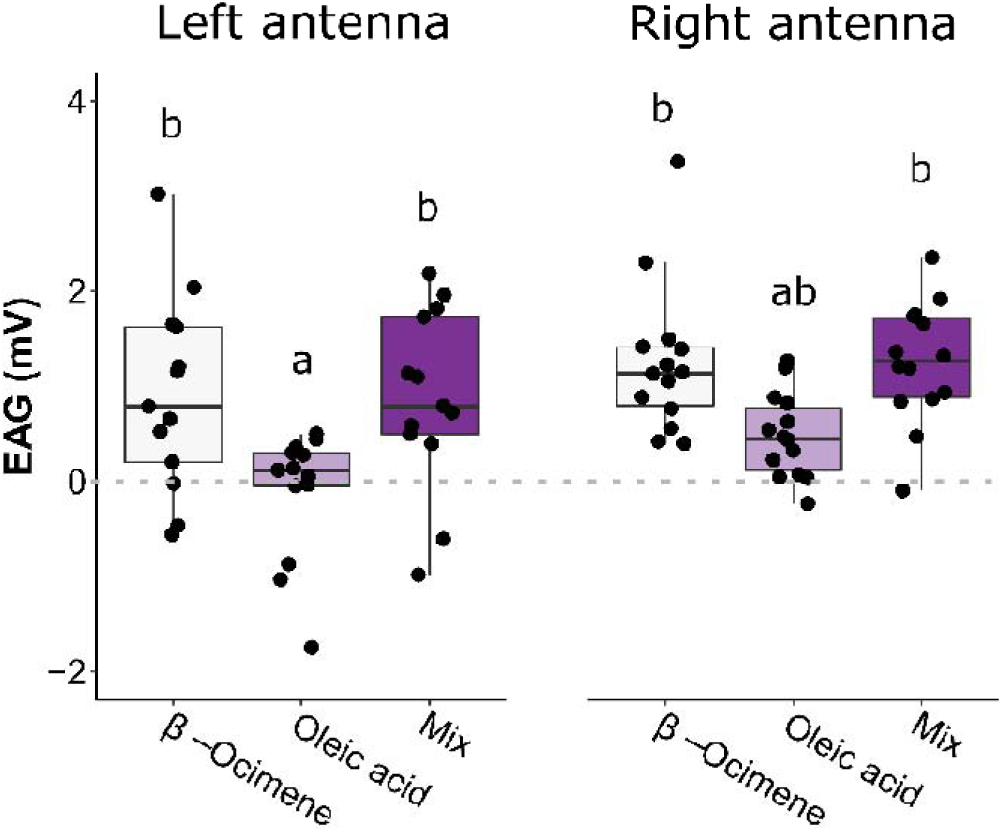
Electroantennography (EAG) responses to odorants. We excised left (N = 13) and right (N = 14) antennae from honey bees in a single highly hygienic colony (score = 95%) and measured the EAG response to neat odorants (Syntech™ CS-55) at hive-realistic temperatures (around 33oC). The EAG response represents blank-subtracted odorant stimuli. We found a significant effect of odorant (two-way ANOVA; levels: side, odorant; F = 12.4, p = 2.3e-5), and letters indicate groups that are significantly different from one another (Tukey HSD p < 0.05). Boxes depict the interquartile range (IQR) and whiskers span 1.5*IQR.

Recently, several antennal protein biomarkers for hygienic behavior have emerged, including two odorant binding proteins (OBP16 and OBP18) which are up-regulated in hygienic bees. To test if β-ocimene and oleic acid are strong ligands for these proteins, we performed *in vitro* binding assays with OBP16 and OBP18. Like our front-way behavioral assays, we used hexane as the odorant negative control and we included phenethyl acetate despite the surprising outcomes of behavioral tests. We found that of the four tested odorants, hexane and phenethyl acetate consistently had the lowest binding affinity (**Figure 6, Table S3**), which mirrors the behavioral response to these compounds (**Figure 2B**). β-ocimene bound OBP16 strongly, but not OBP18. Oleic acid, however, bound both OBPs strongly, with OBP18 being the strongest.

**Figure 6.**
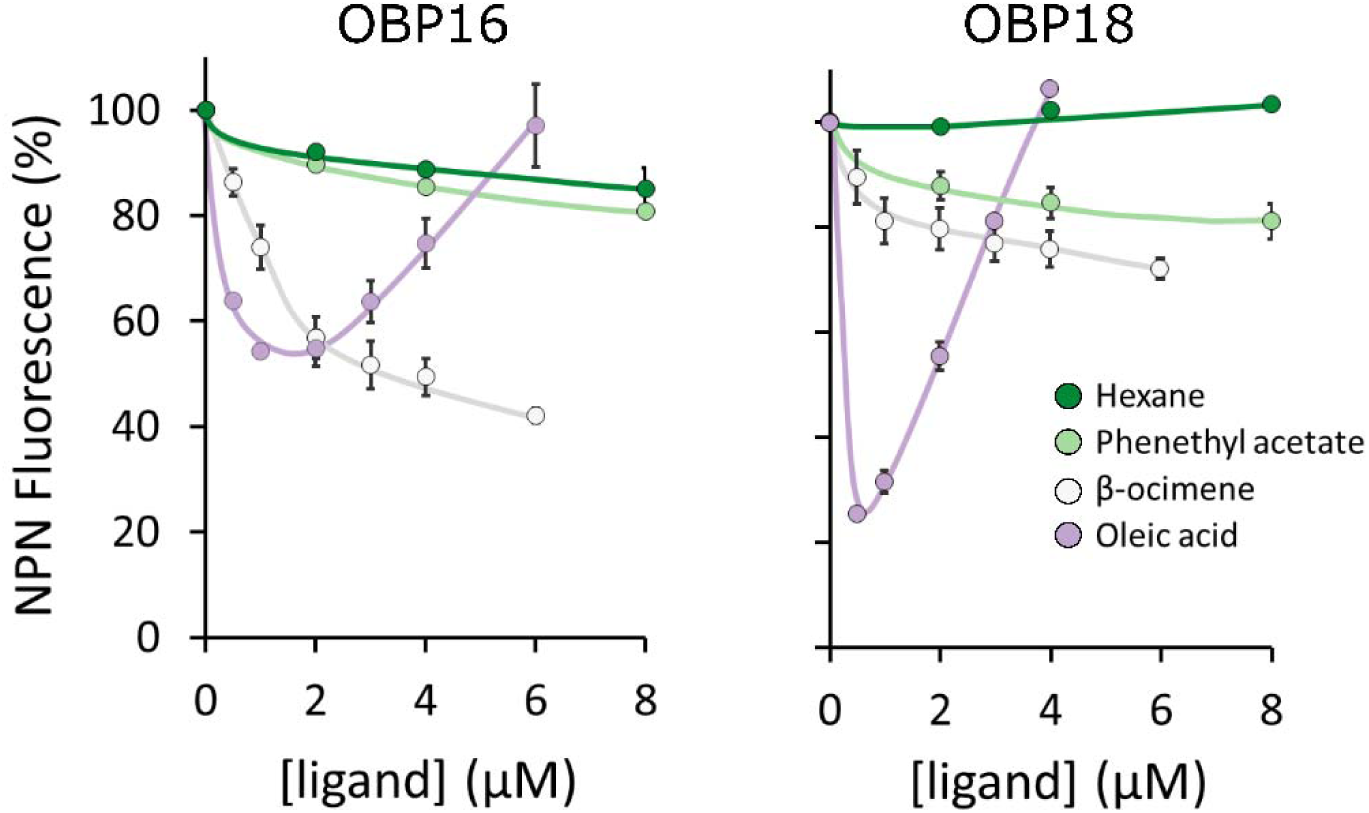
Affinity curves for OBP16 and OBP18. We used an NPN (N-Phenyl-1-naphthylamine) competitive binding assay to measure affinities of β-ocimene, oleic acid, phenethyl acetate, and hexane (negative control). Assays were performed in technical duplicate with 2 µM protein and 2 µM NPN in all cases. Lower NPN fluorescence intensity indicates stronger ligand binding. The high NPN fluorescence intensity for the high oleic acid concentrations is due to the formation of micelles at higher concentrations of the ligands55,56. A 1% solution of β-ocimene, oleic acid, phenethyl acetate, and hexane corresponds to approximately 60 mM, 32 mM, and 63 mM, and 76 mM, respectively. Error bars are standard error of the mean.

## Discussion

Hygienic behavior has been studied in honey bees since at least the 1960s15, but our knowledge of the molecular mechanism behind it is incomplete. In the present work, we investigate two candidate hygienic behavior inducers that are emitted from freeze-killed brood – β-ocimene (a co-opted pheromone emitted by brood and queens^43,44,46-48^) and oleic acid (a well-known necromone and necrophobic compound in other arthropods^1,6,9,10,29-34^) – using *in vivo, ex vivo*, and *in vitro* techniques. We demonstrated 1) that treating brood with the odorants is sufficient to induce hygienic behavior in realistic behavioral assays (**Figure 2B** and **3B**), 2) despite being a viscous compound, oleic acid can stimulate nerve depolarizations worker antennae at hive temperatures (**Figure 5**), and 3) oleic acid and β-ocimene have high affinities to odorant binding proteins that are upregulated in hygienic honey bees (**Figure 6**). Although these specific compounds are from freeze-killed brood and do not extrapolate to all brood diseases, it is a relevant model with which to investigate some molecular interactions governing this complex behavioral process. This is not the first time that a brood pheromone has been implicated in social immunity; Mondet *et al.*^49^ found that *Varroa*-infested brood produced elevated levels of brood ester pheromone. Other researchers have found that oleic acid is both contained and emitted by *Varroa*, on top of it being generally associated with insect death^50-53^.

β-ocimene and oleic acid have very different chemical properties: β-ocimene is a volatile alkene (boiling point: 65-66oC) and oleic acid is a viscous, mono-unsaturated carboxylic acid (boiling point: 360oC). Both are emitted more strongly in freeze-killed honey bee brood compared to live brood^40^, but based on their differences in volatility, we expect them to permeate the brood cell cap at different rates. In a biologically relevant scenario, this spatial diffusion should be necessary for adult workers to detect odorant signals evolving under the cap. Since the odorant blend induces brood removal most consistently in the back-way odorant assays (**Figure 3B**), but not the front-way odorant assays (**Figure 2B**), we suggest that β-ocimene and oleic acid may be acting in a cooperative manner when they have to diffuse through the cap. Since our electroantennogram recordings show that there is no synergistic effect at the level of antennal odorant detection, we suggest they could instead be cooperating via volatility mechanics. For example, a potential mechanism is that β-ocimene diffuses rapidly and attracts worker visits (as it is already known to do for larval feeding48) and after subsequent cell inspection, oleic acid acts as the determinant death cue that stimulates brood removal. In the front-way odorant assay, however, the workers are in constant contact with the odorants (since there is no cap acting as a barrier); therefore, oleic acid is readily detectable even in the absence of an attractant.

The back-way odorant assay we describe here is the most biologically relevant assay for testing different odorants’ abilities to induce hygienic behavior. Unlike other behavioral assays where cells are either uncapped (as in our front-way odorant assay) or filled with odorant-impregnated brood dummies^27,28^, this assay fully maintains comb integrity and allows the workers to perform the complete behavior (uncapping and removal). While the odorant blend was the most consistently high inducer of brood removal, oleic acid alone also induced significant brood removal for young (pre-pupal) brood, but not post-pupal brood (**Figure 3B**). Based on our analysis of the background brood odorant profile, this could be because of naturally released β-ocimene (**Figure 4**) interfering with the synthetic odorant treatments. Since the younger brood emitted significantly more natural β-ocimene compared to the older brood, the young brood treated with oleic acid was, in a way, also a blend, which could explain why this treatment induced similar removal to the synthetic blend for the pre-pupal brood but not post-pupal. Very few pre-pupal β-ocimene-treated brood were removed (28%), which is consistent with young brood emitting their own β-ocimene already. Post-pupal β-ocimene-treated brood, which emit very little natural β-ocimene, were removed at higher rates (54%), although this was not statistically significant (p = 0.10).

With a large body of research showing that olfaction is important for hygienic behavior, combined with two odorant binding proteins (OBP16 and OBP18) emerging as protein biomarkers for hygienic behavior, a tempting hypothesis is that the OBPs are aiding the detection of odorants associated with disease or death. After showing that bees with higher hygienicity remove more β-ocimene- and oleic acid-treated brood compared to bees with lower hygienicity, we performed ligand binding assays with OBP16 and OBP18 to test if the OBPs linked to hygienic behavior have a high affinity to these odorants (**Figure 6**). Interestingly, both hexane and phenethyl acetate had a low affinity to both OBPs, which is consistent with both odorants being poor inducers of hygienic behavior (as demonstrated in our behavioral assays; **Figure 2B**). β-ocimene, however, displayed strong affinity for OBP16. Oleic acid was a strong ligand for both OBPs, and bound OBP18 the strongest of all those we tested. Since β-ocimene and the odorant blend induced significantly higher antennal nerve depolarizations than oleic acid (**Figure 5**), this suggests that either the worker bees must be very close to the emanating cell (or possibly even contacting the source) to sense it, or the odorant treatment induces the brood to emit a different, more volatile signal.

Swanson *et al.*^27^ originally identified phenethyl acetate as a strong hygienic behavior-inducing compound emitted from chalkbrood-infected larvae; however, in our experiments, we found that it induces similar levels of hygienic behavior relative to the negative control in both the diluted (p = 0.99) and neat (p = 0.97) odorant treatments, which is less than both oleic acid and β-ocimene. In fact, Swanson *et al.* found that phenethyl acetate induced 40-100% brood removal using just 5% of the odorant amount we used. One reason why we did not observe high phenethyl acetate removal rates could simply be because the colonies used by Swanson *et al.*^27^ were from a genetic lineage that was more sensitive to chalkbrood odorants than ours. Indeed, the two populations of colonies are geographically isolated and are likely adapted to different climates, conditions, and disease challenges. Furthermore, the surprisingly low degree of overlap between differential expression studies comparing hygienic and non-hygienic bees suggests that there are many adaptive routes for bees to become hygienic54. It could simply be that the hygienic bees in Swanson *et al.*^27^ possess different molecular machinery that allows them to be sensitive to different disease odorants than the colonies used in the present study.

Based on our data, we cannot yet rule out the possibility that some of the behavioral response toward odorant-treated brood was a result of toxicity of the odorant itself. In an acute toxicity assay we found that 1 µl of oleic acid was sufficient to cause contact toxicity when dispensed on the abdomen of pupae, halting the development of 40% of treated individuals and inducing a prophenoloxidase immune response after 2.5 d (Table S4). However, 100% of hexane- and β-ocimene-treated brood developed normally, and the odorant blend caused a response midway between these extremes. While the pattern of removal rates for some of the front-way assay treatments is similar to the toxicity treatments, these differences were not significant. Furthermore, the removal rates for the back-way assays – particularly those post-pupation, which is directly comparable to the toxicity assays in terms of application site and brood age – does not mirror the outcome of the toxicity assay. That being said, the only toxicity outcomes we measured was the prophenoloxidase response and developmental delay. There could be other cues that odorant contact stimulates the brood to emit, which we did not measure. In addition, we only investigated abdominal contact toxicity, which is the application site for the back-way assays, whereas in the front-way assays, we applied odorants to the head, which could yield a different response. Other limitations include that the toxicity outcome was measured after 2.5 d, when other developmental effects could take longer to appear. We note, however, that 2.5 d is much longer than the duration of any of our behavioral assays here. In addition, we tested only pupae in the toxicity assay, and not 5th instar larvae or prepupae, which could respond differently to the odorants. These are all important caveats to this work, and warrant further investigation.

One way these concerns can be addressed in the future is by developing an assay utilizing brood dummies instead of real brood to eliminate the brood effect. Swanson *et al.*^27^ developed a similar assay using brood ester pheromone- and odorant-impregnated paraffin brood dummies in open cells, measuring cell capping (non-hygienic activity) and capping refrainment (hygienic activity) as a proxy for hygienic behavior, since worker bees cannot physically remove the paraffin brood dummies from the cells. This eliminates the brood effect, but has the caveat that leaving a cell uncapped is not the same as performing hygienic behavior. In our front-way experiments, we noticed that cells were frequently left both uncapped and uncannibalized – an outcome which would count as hygienic activity if using paraffin brood dummies. We are currently developing a broodless hygienic test that still allows the object to be removed, for example, by removing developing brood through the back of a Jenter™ set and replacing it with a small odorant-treated object (e.g. a ball of paper or cotton).

On one hand, our 100% odorant treatments (1 µl) could be criticized as not being biologically relevant because the signal is too strong; however, this may work to our advantage to overcome the brood effect. By using such a strong odorant signal in the front-way assays, and measuring the behavior response after a short period of time (3 h, compared to 24 h for the standard freeze-killed brood assay to measure hygienicity), this should a) minimize the amount of time the brood has to produce a strong response, and b) the experimental treatment should be the dominant signal. For the back-way assays, a longer incubation period (20 h) was utilized since in preliminary tests the behavioral response after 3 h was too low to be useful. This means that there was more time for a potential brood effect to evolve, which may have impacted our results.

Regardless of the caveats to the behavioral assays, the ligand binding assays provide a clear picture: β-ocimene and oleic acid each strongly bind at least one of the two odorant binding proteins (OBP16 and OBP18) whose expression is more strongly correlated with hygienic behavior^23,35^. While oleic acid produces high fluorescence intensities (which normally indicates weak binding) at higher ligand concentrations (*i.e.* > 1 µM), this is a well-known phenomenon for amphipathic ligands^55,56^. The very low fluorescence intensity < 1 µM indicates that it is indeed a strong ligand for OBP^18^, which agrees with previous binding assays35. Conversely, the two odorants which induced low rates of hygienic behavior in our assays also were poor ligands for these OBPs. Therefore, the results of this *in vitro* binding assay can explain the behavioral observations surprisingly well. Despite this evidence, it’s difficult to know how well the OBP and ligand concentrations reflect reality. For example, the absolute concentration of OBPs in the hemolymph of honey bee antennae is currently unknown, as is the effective ligand concentration at the antennal pore (the interface between the hemolymph and the surrounding air). While a 1% solution of β-ocimene corresponds to approximately a 60 mM solution, which is much higher than the concentrations in the ligand binding assays (<10 µM), with volatility mechanics and spatial diffusion, the airborne concentration is likely much lower (but unknown).

In summary, this data suggests that oleic acid and β-ocimene induce brood removal in honey bees. Bees with higher hygienicity respond to the odorants more strongly than bees with lower hygienicity, and the blend induces brood removal most consistently in the most biologically realistic brood removal assay. Despite being non-volatile, oleic acid appears to be detectable even beneath a brood cell cap; however, it’s possible that the bees are detecting the brood’s reaction to the odorant rather than the odorant alone. Our electrophysiology tests show that oleic acid only marginally stimulates antennal nerve responses in environmental conditions similar to those inside a hive, suggesting that if they are detecting the odorant alone, extremely close proximity would be necessary for bees to detect it. Both odorants are strong ligands for at least one of the OBPs linked to hygienic behavior, whereas hexane and phenethyl acetate (which induced the lowest levels of hygienic behavior) are weak ligands for both OBPs. These molecular interactions between the odorant ligands and the OBPs mirror the results of our behavioral assay surprisingly well. Furthermore, oleic acid elicits necrophoretic and necrophobic behavior across phylum Arthropoda^1,6,9,10,29-34^, and these data piece its activity in honey bees into the phylogenetic puzzle. To the best of our knowledge, our data shows for the first time that this ‘death cue’ function is evolutionarily conserved in honey bees, and that oleic acid may be working in concert with β-ocimene as an attractant. Future experiments will be necessary to eliminate the possibility of an odorant-induced brood effect contributing to these results.

## Methods

### Honey bee colonies and hygienic testing

We kept honey bee colonies at four separate apiaries in Greater Vancouver, Canada, and performed hygienic testing as previously described^23^. Briefly, for each test, polyvinyl chloride pipes (5 cm inner diameter, ∼25 cm length) were pressed into capped brood comb in two areas containing white-eyed to red-eyed pupae, then filled with approximately 250 ml of liquid nitrogen to freeze. Frames were returned to the colony and assessed 24 h later for percent removal of the frozen brood cells. One week later, the test was repeated, and the average of the two tests (four 5 cm brood patches in total) yielded the FKB score. All hygienic testing, sampling and odorant assays were completed during the summer of 2017.

### Front-way odorant assays

To perform the front-way odorant assays, we retrieved two brood frames from each colony, uncapped patches of brood with tweezers and dispensed 1 µl of odorant treatments onto the exposed brood (**Figure 1A**). Wax caps were not replaced after odorant addition. We tested the odorants β-ocimene, oleic acid, a 1:1 v/v blend of the two, phenethyl acetate (positive control), and hexane (negative control) at concentrations of 100% and 1% (v/v in hexane). Phenethyl acetate was not included in the blend because it is not known to co-occur with the other odorants (phenethyl acetate is from chalkbrood, while β-ocimene and oleic acid are associated with freeze-killed brood). For each odorant and concentration, we performed two technical replicates (2 patches of 30 brood cells each, one on each frame). We tested the different concentrations on different days. After treating the brood patches with odorants, we photographed, traced, and labelled each patch on a transparency and replaced the brood frame in the hive. After 3 h, we returned to the hive and recorded the number of brood cells that were cannibalized and partially cannibalized (cumulatively yielding the number ‘removed‘) or recapped.

Brood patches were composed of variable developmental stages (mostly prepupae and pupae, but some 5th instar larvae; Table S5), so we used the photographs from pre- and post-incubation to assess the fraction of each developmental stage that were removed and/or recapped by the workers. With a clear anterior view, the prepupae can be distinguished from 5th instar larvae based on their upright, elongated body and a ‘crook-neck’ appearance. Due to variable patch composition, we did not obtain the same number of biological replicates for every developmental stage and odorant (see Table 1 for complete replicate information for each stage and odorant concentration). Data for 5th instar larvae are not shown because too few patches contained them to reliably test if there was a differential response to larvae (they made up < 10% of tested brood cells overall). This is because the time between cell capping and transforming to a prepupa is very short – in the order of hours – so catching this stage in a naturally laid comb is infrequent. These sparse data were therefore excluded from subsequent analyses. In a preliminary test, brood removal data were analyzed with a four-factor ANOVA (levels: dose, odorant, age, hygienicity) followed by a Tukey HSD to determine if there was an effect of age between prepupae and pupae. Since there was no significant effect of age alone (F = 0.87; p = 0.36) nor in combination with any other factors (odorant*age: p = 0.61, dose*age: p = 0.15, hygienicity*age: p = 0.79, odorant*dose*age: p = 0.58, odorant*hygienicity*age: p = 0.73, dose*hygienicity*age: p = 0.17, odorant*dose*hygienicity*age: p = 0.71), we pooled the pupa and prepupa data for subsequent analyses. All statistical analyses were performed in R unless otherwise specified.

In a second preliminary experiment, we confirmed that there was no effect of patch proximity in the front-way odorant assay. We varied proximity by testing two patches of brood per colony that were either separated by a single capped cell-width on the same side of a frame (’near‘), or on different frames with two brood frames located between them (’far,’ N = 5 colonies each). One microliter of oleic acid (the least volatile odorant tested) or β-ocimene (the most volatile odorant tested) was added to the cells of each patch. The data were analyzed with a two-factor ANOVA (levels: proximity, odorant).

To assess the relationship between hygienicity and odorant-treated brood removal, we performed the front-way odorant assay on 10 colonies (and two technical replicates per colony, which were averaged to produce one biological replicate) with varying hygienic score (39% to 100%). We grouped the colonies into N = 5 with higher hygienicity (scoring > 80%), and N = 5 with lower hygienicity (scoring < 80%) (**Figure 2A**). As before, we removed the larval cells from the analysis (∼10% overall) and since we previously determined that there was no effect of brood age between prepupae and pupae in the front-way assay, we did not distinguish between these stages statistically. These data were analyzed using a three-factor ANOVA (levels: dose, odorant, hygienicity) followed by a Tukey HSD post hoc test. Brood recapping data was derived from the same assays (N = 9 for each odorant (data was unavailable for one colony)) using a one-way ANOVA (level: odorant).

### Back-way odorant assays

To test the effects of β-ocimene, oleic acid and their 1:1 v/v blend in a more biologically realistic scenario, we developed the back-way odorant assay (**Figure 3A**). This assay adapts artificial comb cages of the Jenter™ queen rearing system to instead rear worker brood *in situ*. The Jenter™ set features removable plastic plugs from the rear of the comb – usually used to harvest eggs/larvae for queen rearing – which provide convenient access points for odorant addition without damaging the wax brood cell caps or the brood itself.

We conditioned the Jenter™ comb cages by placing them in a colony for several days, allowing the bees to draw out full-height comb cells. We then caged the queens and allowed them sufficient time to populate the combs with eggs (typically overnight). We released the queens and allowed the workers to rear the brood *in situ*. Once capped, we inspected the brood via the removable plugs to confirm the developmental stage. Through this small posterior window, 5th instar larvae and prepupae are indistinguishable, but pupae are easily recognized by their clearly developed abdomen and hind feet. This is in contrast to the front-way odorant assay, where 5th instar larvae and prepupae are distinguishable due to the clear anterior view of the head.

We removed the plugs for 9-10 semi-randomly located brood cells (each group of 9-10 cells in a different colony = 1 biological replicate) and dispensed odorants (1 µl of neat solutions) onto the brood through the back of the comb and re-plugged each cell. The number of brood in these patches is smaller than for the front-way odorant assays because the size of the Jenter™ cage limits the total brood area. We traced a map of the odorant-treated cells and placed the combs in colonies for 20 h to allow workers to detect the odorant signals through the cap and respond. We performed five biological replicates (*i.e.* repeated the test in five colonies) for each odorant and developmental stage (pre-pupation and post-pupation). Since the 5th instar larvae and pre-pupae are indistinguishable (as described above), the ‘pre-pupation’ group contains both stages. After incubation, we removed the comb and counted the number of brood cells from each odorant treatment that were removed and/or partially cannibalized. Removal data was analyzed as described above except we used a two-factor ANOVA (levels: odorant, age). Due to spontaneous re-queening events and subsequent worker turn-over, the hygienic scores are not known for all of the colonies in this experiment.

### Gas chromatography mass spectrometry (GC-MS)

We performed GC-MS on extracts from larvae, prepupae and pupae to detect differences in their natural odorant profiles. Here, the three stages are distinguishable because by removing the brood from the cell, we can clearly differentiate the features of a prepupae compared to a 5th instar larva (the elongated body and ‘crook-neck’ appearance). We collected capped 5th instar larvae, prepupae and pupae from five different colonies and performed solid-phase micro-extraction (SPME) GC-MS as well as cuticle hexane wash GC-MS as previously described40. Briefly, for the SPME analysis, we sealed individual brood in 10 mL glass vials (having previously confirmed that this is sufficient air to prevent suffocation) for 24 h prior to GC-MS analysis to allow volatiles to equilibrate in the headspace. We analyzed N = 5 brood for each stage, with all 5 coming from different colonies, in a sample order randomized by colony and stage to avoid batch effects. The extracted compounds were analyzed by low-resolution GC-MS (Agilent 789011A/597511C Inert XL MSD) with a DB-wax analytical column (J&W 122–7032) and a 45 min temperature gradient spanning from 50oC to 230oC. For the hexane wash analysis, we soaked individual brood (also N = 5 brood for each developmental stage, from 5 different colonies) in 300 µl of HPLC-grade hexane for 5 min with gentle agitation. We injected 1 µl of each sample on to the analytical column (same as above) connected to a Agilent 6890⍰N/5975⍰C Inert XL MSD mass spectrometer. The temperature gradient spanned the same temperatures but was 30 min long.

Spectral data was searched using Mass Hunter Qualitative Analysis software (vB.06.00) and the Wiley Chemical Compound Library (W9N08.L). Since the automatic integration algorithm within Mass Hunter often applies erroneous peak baselines, peak areas were integrated manually. Only peaks with apex intensities exceeding 4,000 cts were integrated, since less intense peaks rarely yielded confident spectral matches to known compounds. Raw data is available for download at http://github.com/AlisonMcAfee/test. Peak areas were log10 transformed and compound profiles were compared between developmental stages using a one-way ANOVA followed by a Benjamini-Hochberg correction (5% FDR) performed in Perseus (v1.5.5.3). We confirmed the identity of β-ocimene, isopropanol, and 2-pentanone against synthetic standards (Sigma).

### Electroantennography (EAG) recordings

We obtained EAG recordings on bees’ antennae collected from a single highly hygienic colony (freeze-killed brood score = 95%) maintained at the University of British Columbia. For *ex vivo* EAG analysis, we sampled adult nurses from an open brood frame and kept them in a humid incubator (35°C) with access to sucrose water (1:1) until antennal excision. Immediately prior to EAG testing, either the left or right antenna was removed from individual bees (according to *a priori* randomization) by cutting at the base of the scape. We trimmed the last flagellum segment with dissection scissors, then connected the ends to recording electrodes via glass capillary tubes filled with insect saline solution (210 mM NaCl, 3.1 mM KCl, 10 mM CaCl_2_, 2.1 mM NaCO_3_, and 0.1 mM NaH_2_ PO_4_, pH 7.2) as previously described57. After data was acquired for the first antenna, the second was collected and the bee was euthanized. In total we acquired data for N = 13 left antennae and N = 14 right antennae.

During EAG acquisition, we used a Syntech™ CS-55 stimulus controller to continuously pass humidified air over the antenna and to deliver 1 s pulses of odorized air. To produce odorized air, we cut 1 cm2 slips of No. 1 Whatman filter paper and inserted them into glass Pasteur pipette cartridges. We heated the cartridges to 37°C using a flexible chromatography column heater, at which time we dispensed onto the filter paper 5 µl of distilled water (blank), β-ocimene, oleic acid, or a 1:1 v/v blend of β-ocimene and oleic acid (mix). After allowing 30 s of initial evaporation and slight cooling for the cartridge to reach approximately 33oC, we aimed away from the antenna and passed a 1 s burst of room-temperature air through the pipette before stimulating the antennae with the odorants. We then exposed the antennae to a set of 3 consecutive 1 s bursts for each odorant in a randomly-determined order. Between 0.5 and 1 min was allowed between each presentation to allow antennal electrical activity to return to baseline. Blank stimuli (also 3 consecutive 1 s bursts each) were performed at two randomly determined times during acquisition. For each antenna, we subtracted the average blank intensity from the odorant EAG intensities, then compared odorant groups with a two-way ANOVA (levels: odorant, side).

### Ligand affinity assays for odorant binding proteins (OBPs)

Recombinant OBP16 and OBP18 were cloned, expressed, and purified exactly as previously described35. Briefly, the OBP genes were PCR amplified from honey bee cDNA and cloned into a PET-5b bacterial expression vector. Plasmids were transformed into BL21(DE3)Rosetta-gami (OBP16) and BL21(DE3)pLysS E. coli strains and protein expression was induced via IPTG. The recombinant proteins were then purified by a series of chromatographic elutions, including anion exchange (DE-52, QFF, or Mono-Q) and gel filtration (Sephacryl-100 or Superose-12) as well as other standard purification protocols^56,58^.

We then used an NPN (N-Phenyl-1-naphthylamine) competitive binding assay to measure relative affinities of β-ocimene, oleic acid, phenethyl acetate, and hexane (negative control). Binding assays were also conducted as previously described(31), except they were performed in technical duplicate with 2 µM protein, 2 µM NPN, and between 0 and 8 µM of hexane and phenethyl acetate or between 0 and 6 µM of β-ocimene and oleic acid. Dissociation constants of the ligands were calculated from the corresponding IC_50_ values (concentrations of ligands halving the initial fluorescence value of 1-NPN), using the equation: KD = [IC_50_]/(1 + [1 - NPN]/K(1 - NPN)) where [1-NPN] is the free concentration of 1-NPN and K(1 - NPN) is the dissociation constant of the complex protein/[1-NPN].

### Odorant toxicity assays

To test the toxicity of the odorants, we retrieved 60 purple-eyed, white body pupae and applied 1 µl of neat odorant (phenethyl acetate was not included) to the dorsal abdominal area (n = 15 each). We placed the pupae in tissue-lined petri dishes and incubated them at 33oC for 2.5 d. We then scored the pupae for whether their development was halted (*i.e.* their cuticle did not begin to brown or harden and their eye pigment did not change colour) and whether a prophenoloxidase response had initiated (*i.e.* the dorsal abdominal region became black). All pupae with halted development also had a prophenoloxidase response.

### Data availability

The datasets generated during and/or analysed during the current study are available from the corresponding author on reasonable request. The raw GC-MS data is available for download at http://www.github.com/AlisonMcAfee/test.

## Acknowledgements

We would like to thank Deborah Tin Tun and Liam Brownrigg for allowing us to access their honey bee colonies for some of our experiments.

## Author contributions

AM wrote the first draft of the manuscript, designed the experiments and collected behavioral data with the help of AC, HH, and LJF. YGK performed the EAG experiments, with assistance from TFC. II and PP performed the *In vitro* binding assays. LLM acquired the GC-MS data.

## Additional information

The authors declare no competing interests.

## References

1 Sun, Q. & Zhou, X. Corpse management in social insects. Int J Biol Sci 9, 313–321, doi:10.7150/ijbs.5781 (2013).

2 Cremer, S., Armitage, S. A. & Schmid-Hempel, P. Social immunity. Curr Biol 17, R693–702, doi:10.1016/j.cub.2007.06.008 (2007).

3 Wilson-Rich, N., Spivak, M., Fefferman, N. & Starks, P. T. Genetic, individual, and group facilitation of disease resistance in insect societies. Annu Rev Entomol 54, 405–423, doi:10.1146/annurev.ento.53.103106.093301 (2009).

4 Wilson, E., Durlach, N. & Roth, L. Chemical releasers of necrophoric behavior in ants. Psyche 65, 108–114 (1958).

5 Haskins, C. P. & Haskins, E. F. Notes on necrophoric behavior in the archaic ant Myrmecia vindex (Formicidae: Myrmeciinae). Psyche 81, 258–267 (1974).

6 Gordon, D. M. Dependence of necrophoric response to oleic acid on social context in the ant, Pogonomyrmex badius. Journal of chemical ecology 9, 105–111 (1983).

7 Howard, D. F. & schinkel, W. R. Aspects of necrophoric behavior in the red imported fire ant, Solenopsis invicta. Behaviour 56, 157–178 (1976).

8 Julian, G. E. & Cahan, S. Undertaking specialization in the desert leaf-cutter ant Acromyrmex versicolor. Animal Behaviour 58, 437–442 (1999).

9 Chouvenc, T., Robert, A., Sémon, E. & Bordereau, C. Burial behaviour by dealates of the termite Pseudacanthotermes spiniger (Termitidae, Macrotermitinae) induced by chemical signals from termite corpses. Insectes sociaux 59, 119–125 (2012).

10 Ulyshen, M. D. & Shelton, T. G. Evidence of cue synergism in termite corpse response behavior. Naturwissenschaften 99, 89–93 (2012).

11 Sun, Q., Haynes, K. F. & Zhou, X. Differential undertaking response of a lower termite to congeneric and conspecific corpses. Sci Rep 3, 1650, doi:10.1038/srep01650 (2013).

12 Le Conte, Y. et al. Social immunity in honeybees (Apis mellifera): transcriptome analysis of varroa-hygienic behaviour. Insect Mol Biol 20, 399–408, doi:10.1111/j.1365-2583.2011.01074.x (2011).

13 Spivak M. & Gilliam, M. Hygienic behaviour of honey bees and its application for control of brood diseases and Varroa: Part II. Studies on hygienic behaviour since the Rothenbuhler era. Bee world 79, 169–186 (1998).

14 Spivak, M. & Gilliam, M. Hygienic behaviour of honey bees and its application for control of brood diseases and varroa: Part I. Hygienic behaviour and resistance to American foulbrood. Bee World 79, 124–134 (1998).

15 Rothenbuhler, W. C. Behavior genetics of nest cleaning in honey bees. IV. Responses of F 1 and backcross generations to disease-killed brood. American Zoologist 4, 111–123 (1964).

16 Boecking, O. & Drescher, W. The removal response ofApis mellifera L. colonies to brood in wax and plastic cells after artificial and natural infestation withVarroa jacobsoni Oud. and to freeze-killed brood. Experimental and Applied Acarology 16, 321–329 (1992).

17 Gilliam, M., Taber III, S. & Richardson, G. V. Hygienic behavior of honey bees in relation to chalkbrood disease. Apidologie 14, 29–39 (1983).

18 Palacio, M. A., Rodriguez, E., Goncalves, L., Bedascarrasbure, E. & Spivak, M. Hygienic behaviors of honey bees in response to brood experimentally pin-killed or infected with Ascosphaera apis. Apidologie 41, 602–612 (2010).

19 Spivak, M. & Reuter, G. Resistance to American foulbrood disease by honey bee colonies Apis mellifera bred for hygienic behavior. Apidologie 32, 555–565 (2001).

20 Ibrahim, A. & Spivak, M. The relationship between hygienic behavior and suppression of mite reproduction as honey bee (Apis mellifera) mechanisms of resistance to Varroa destructor. Apidologie 37, 31–40 (2006).

21 Spivak, M. Honey bee hygienic behavior and defense against Varroa jacobsoni. Apidologie 27, 245–260 (1996).

22 Bixby, M. et al. A Bio-Economic Case Study of Canadian Honey Bee (Hymenoptera: Apidae) Colonies: Marker-Assisted Selection (MAS) in Queen Breeding Affects Beekeeper Profits. J Econ Entomol 110, 816–825, doi:10.1093/jee/tox077 (2017).

23 Guarna, M. M. et al. Peptide biomarkers used for the selective breeding of a complex polygenic trait in honey bees. Sci Rep 7, 8381, doi:10.1038/s41598-017-08464-2 (2017).

24 Gramacho, K. P. & Spivak, M. Differences in olfactory sensitivity and behavioral responses among honey bees bred for hygienic behavior. Behavioral Ecology and Sociobiology 54, 472–479 (2003).

25 Masterman, R., Ross, R., Mesce, K. & Spivak, M. Olfactory and behavioral response thresholds to odors of diseased brood differ between hygienic and non-hygienic honey bees (Apis mellifera L.). Journal of Comparative Physiology A 187, 441–452 (2001).

26 Spivak, M., Masterman, R., Ross, R. & Mesce, K. A. Hygienic behavior in the honey bee (Apis mellifera L.) and the modulatory role of octopamine. Journal of neurobiology 55, 341–354 (2003).

27 Swanson, J. A. et al. Odorants that induce hygienic behavior in honeybees: identification of volatile compounds in chalkbrood-infected honeybee larvae. Journal of chemical ecology 35, 1108–1116 (2009).

28 Nazzi, F., Della Vedova, G. & D’Agaro, M. A semiochemical from brood cells infested by Varroa destructor triggers hygienic behaviour in Apis mellifera. Apidologie 35, 65–70 (2004).

29 Yao, M. et al. The Ancient Chemistry of Avoiding Risks of Predation and Disease. Evolutionary Biology 36, 267–281, doi:10.1007/s11692-009-9069-4 (2009).

30 Aksenov, V. & David Rollo, C. Necromone Death Cues and Risk Avoidance by the Cricket Acheta domesticus: Effects of Sex and Duration of Exposure. Journal of Insect Behavior 30, 259–272, doi:10.1007/s10905-017-9612-6 (2017).

31 Qiu, H. L. et al. Differential necrophoric behaviour of the ant Solenopsis invicta towards fungal-infected corpses of workers and pupae. Bull Entomol Res 105, 607–614, doi:10.1017/S0007485315000528 (2015).

32 Akino, T. & Yamaoka, R. Origin of Oleic Acid, Corpse Recognition Signal in the Ant, Formica japonica MOTSCHLSKY (Hymenoptera: Formicidae). Japanese journal of applied entomology and zoology 40, 265–271, doi:10.1303/jjaez.40.265 (1996).

33 Abbott, K. R. Bumblebees avoid flowers containing evidence of past predation events. Canadian Journal of Zoology 84, 1240–1247, doi:10.1139/z06-117 (2006).

34 Rollo, C. D., Czvzewska, E. & Borden, J. H. Fatty acid necromones for cockroaches. Naturwissenschaften 81, 409–410, doi:10.1007/bf01132695 (1994).

35 Guarna, M. M. et al. A search for protein biomarkers links olfactory signal transduction to social immunity. BMC Genomics 16, 63, doi:10.1186/s12864-014-1193-6 (2015).

36 Hu, H. et al. Proteome analysis of the hemolymph, mushroom body, and antenna provides novel insight into honeybee resistance against Varroa infestation. Journal of Proteome Research (2016).

37 Mondet, F. et al. Antennae hold a key to Varroa-sensitive hygiene behaviour in honey bees. Scientific reports 5, 10454 (2015).

38 Parker, R. et al. Correlation of proteome-wide changes with social immunity behaviors provides insight into resistance to the parasitic mite, Varroa destructor, in the honey bee (Apis mellifera). Genome biology 13, 1 (2012).

39 Forêt, S. & Maleszka, R. Function and evolution of a gene family encoding odorant binding-like proteins in a social insect, the honey bee (Apis mellifera). Genome Res 16, 1404–1413, doi:10.1101/gr.5075706 (2006).

40 McAfee, A., Collins, T. F., Madilao, L. L. & Foster, L. J. Odorant cues linked to social immunity induce lateralized antenna stimulation in honey bees (Apis mellifera L.). Sci Rep 7, 46171, doi:10.1038/srep46171 (2017).

41 Spivak, M. & Downey, D. L. Field assays for hygienic behavior in honey bees (Hymenoptera: Apidae). Journal of economic entomology 91, 64–70 (1998).

42 He, X. J. et al. Starving honey bee (Apis mellifera) larvae signal pheromonally to worker bees. Scientific reports 6 (2016).

43 Maisonnasse, A. et al. A scientific note on E-$\ beta $-ocimene, a new volatile primer pheromone that inhibits worker ovary development in honey bees. Apidologie 40, 562–564 (2009).

44 Maisonnasse, A., Lenoir, J. C., Beslay, D., Crauser, D. & Le Conte, Y. E-β-ocimene, a volatile brood pheromone involved in social regulation in the honey bee colony (Apis mellifera). PLoS One 5, e13531, doi:10.1371/journal.pone.0013531 (2010).

45 Traynor, K. S., Le Conte, Y. & Page, R. E. Age matters: pheromone profiles of larvae differentially influence foraging behaviour in the honeybee, Apis mellifera. Animal Behaviour 99, 1–8 (2015).

46 Ma, R., Mueller, U. G. & Rangel, J. Assessing the role of β-ocimene in regulating foraging behavior of the honey bee, Apis mellifera. Apidologie 47, 135–144, doi:10.1007/s13592-015-0382-x (2016).

47 Gilley, D. C., Degrandi-Hoffman, G. & Hooper, J. E. Volatile compounds emitted by live European honey bee (Apis mellifera L.) queens. J Insect Physiol 52, 520–527, doi:10.1016/j.jinsphys.2006.01.014 (2006).

48 He, X. J. et al. Starving honey bee (Apis mellifera) larvae signal pheromonally to worker bees. Sci Rep 6, 22359, doi:10.1038/srep22359 (2016).

49 Mondet, F. et al. Specific Cues Associated With Honey Bee Social Defence against Varroa destructor Infested Brood. Scientific reports 6 (2016).

50 Plettner, E., Eliash, N., Singh, N. K., Pinnelli, G. R. & Soroker, V. The chemical ecology of host-parasite interaction as a target of Varroa destructor control agents. Apidologie 48, 78–92 (2017).

51 Martin, C. et al. Potential mechanism for detection by Apis mellifera of the parasitic mite Varroa destructor inside sealed brood cells. Physiological Entomology 27, 175–188 (2002).

52 Zalewski, K., Zaobidna, E. & Żóltowska, K. Fatty acid composition of the parasitic mite Varroa destructor and its host the worker prepupae of Apis mellifera. Physiological Entomology 41, 31– 37 (2016).

53 Dmitryjuk, M., Zalewski, K., Raczkowski, M. & Zoltowska, K. Composition of fatty acids in the Varroa destructor mites and their hosts, Apis mellifera drone-prepupae. Annals of parasitology 61 (2015).

54 Gempe, T., Stach, S., Bienefeld, K., Otte, M. & Beye, M. Behavioral and molecular studies of quantitative differences in hygienic behavior in honeybees. BMC Res Notes 9, 474, doi:10.1186/s13104-016-2269-y (2016).

55 Leal, G. M. & Leal, W. S. Binding of a fluorescence reporter and a ligand to an odorant-binding protein of the yellow fever mosquito, Aedes aegypti. F1000Research 3 (2014).

56 Sun, Y. F. et al. Two odorant-binding proteins mediate the behavioural response of aphids to the alarm pheromone (E)-ß-farnesene and structural analogues. PLoS one 7, e32759 (2012).

57 Olsson, S.B. & Hansson, B. S. Electroantennogram and single sensillum recording in insect antennae. Methods Mol Biol 1068, 157–177, doi:10.1007/978-1-62703-619-1_11 (2013).

58 Ban, L. et al. Chemosensory proteins of Locusta migratoria. Insect Mol Biol 12, 125–134 (2003).

